# Electric field dependent effects of motor cortical TDCS

**DOI:** 10.1101/327361

**Authors:** Ilkka Laakso, Marko Mikkonen, Soichiro Koyama, Daisuke Ito, Tomofumi Yamaguchi, Akimasa Hirata, Satoshi Tanaka

## Abstract

Transcranial direct current stimulation (TDCS) can modulate motor cortical excitability. However, its after-effects are highly variable between individuals. Individual cranial and brain anatomy may contribute to this variability by producing varying electric fields in each subject’s brain. Here we show that these fields are related to excitability changes following anodal TDCS of the primary motor cortex (M1). We found in two experiments (N=28 and N=9) that the after-effects of TDCS were proportional to the individual electric field in M1, calculated using MRI-based models. Individuals with the lowest and highest local electric fields in M1 tended to produce opposite changes in excitability. Furthermore, the effect was field-direction dependent and non-linear with stimulation duration or other experimental parameters. The electric field component pointing into the brain was negatively proportional to the excitability changes following 1 mA 20 min TDCS of right M1 (N=28); the effect was opposite after 1 mA 10 min TDCS of left M1 (N=9). Our results demonstrate that a large part of variability in the after-effects of motor cortical TDCS is due to inter-individual differences in the electric fields. We anticipate that individualized electric field dosimetry could be used to control the neuroplastic effects of TDCS, which is increasingly being explored as a treatment for various neuropsychiatric diseases.

## 1 Introduction

Transcranial direct current stimulation (TDCS) is a widely used non-invasive method capable of eliciting changes in cortical excitability (Priori et al., 1998; Nitsche and Paulus, 2000, 2001). These neuroplastic changes have potential as a treatment for various psychiatric and neurological diseases that involve pathological changes in plasticity (Kuo et al., 2014; Flöel, 2014). Cortical excitability changes induced by TDCS can be most reliably measured in the primary motor cortex (M1) using transcranial magnetic stimulation (TMS) to measure the amplitude of the motor evoked potentials (MEP) (Horvath et al., 2015). In such studies, the responses to TDCS have been found to be highly variable between individuals (Wiethoff et al., 2014; López-Alonso et al., 2014; Chew et al., 2015; Strube et al., 2016; Ammann et al., 2017). The underlying reasons of the variability are still unknown.

The physical agent of TDCS is thought to be the electric field (EF) that is generated in the brain and other tissues when direct current (usually 1-2 mA) is applied through electrodes attached to the scalp. In the brain, the EF is weak, typically less than 1 V/m in strength (Datta et al., 2009; Truong et al., 2013; Opitz et al., 2015; Laakso et al., 2015, 2016). Animal *in vitro* studies have shown that such weak EFs can affect the activity of M1 (Bikson et al., 2004; Fritsch et al., 2010). Long-lasting excitability changes produced by weak EFs may depend on NMDA receptors (Fritsch et al., 2010), which is also supported by electrophysiological studies in humans, where oral intake of NMDA antagonist suppressed the after-effects of TDCS (Nitsche et al., 2003).

We have previously found that there are large differences in the EFs between individuals (Laakso et al., 2015). The differences are due to anatomical factors, such as gyral and sulcal anatomy as well as the thicknesses of the CSF and scalp (Datta et al., 2012; Laakso et al., 2015; Opitz et al., 2015). However, the role of EFs in interindividual variability is still unclear. Are the effects of EF on neural tissue sufficiently similar in each individual so that EFs were useful for predicting the effects of TDCS? If they were, individual EF models could hypothetically be used to reduce variability and control the effects.

Here, we studied whether the EF was related to the after-effects of TDCS. We first performed an exploratory sham-controlled motor cortical TDCS study and individually calculated the EFs in all our subjects. As the effects of TDCS are sensitive to the field direction (Rawji et al., 2018), we analysed all three orthogonal components of the EF. To find which cortical sites are potentially affected by the EF, we decided to use partial least squares (PLS) regression (Geladi and Kowalski, 1986; Wold et al., 2001), which is an effective method for finding relationships between dependent variables (here: MEP amplitude) and a large number of collinear predictor variables (here: EF in the cortex). Compared to other commonly used approaches for feature extraction from imaging data, such as random field theory (Worsley et al., 1996), PLS regression was advantageous because we needed not define a region of interest *a priori,* which would have been arbitrary as we did not know in advance which site in M1 or other regions (Fischer et al., 2017) was affected by TDCS. At the potentially important cortical site, the data were further analysed using linear mixed effects models to investigate the direction and persistence of the effects. A second experiment was conducted to validate the effects found in the main experiment.

## 2 Methods

### 2.1 Subjects

Thirty-seven healthy subjects (10 females and 27 males) participated in the experiments. Twenty-eight subjects (7 females and 21 males; mean age ± SD = 27±6 years) participated in experiment 1, and nine subjects (3 females and 6 males; mean age ± SD = 28±6 years) participated in experiment 2.

The subjects were neurologically healthy and had no family history of epilepsy. All subjects gave informed consent before participating in the experiments. The Human Ethics Committee at the National Institute for Physiological Sciences, Okazaki, Japan, approved experiment 1, and Ethics Committee at Tokyo Bay Rehabilitation Hospital, Chiba, Japan, approved experiment 2. All methods were carried out in accordance with approved institutional guidelines and regulations.

### 2.2 MRI

All subjects participated in MRI scanning. T1- and T2-weighted structural MRI scans of subjects participating in experiment 1 were acquired using a 3.0 T MRI scanner (Verio; Siemens, Ltd., Erlangen, Germany). Subjects participating in experiment 2 were imaged using a 1.5 T MRI scanner (Intera; Philips Healthcare, Ltd., Andover, Netherlands).

### 2.3 Experimental parameters

TDCS (1 mA) was applied using a DC STIMULATOR PLUS (NeuroConn, Germany) in two experiments with two conditions each, which are summarized in Fig. 1. Conditions were separated by a washout period of at least three days. As a measure of cortical excitability, MEPs were elicited using a Magstim 200^2^ magnetic stimulator (Magstim Company, UK). At the beginning of each condition, we determined the resting motor threshold (RMT). RMT was defined as the lowest stimulation intensity required to elicit MEPs of 50 *μ*V peak-to-peak amplitude in five of ten trials (Rossini et al., 1999).

**Figure 1:**
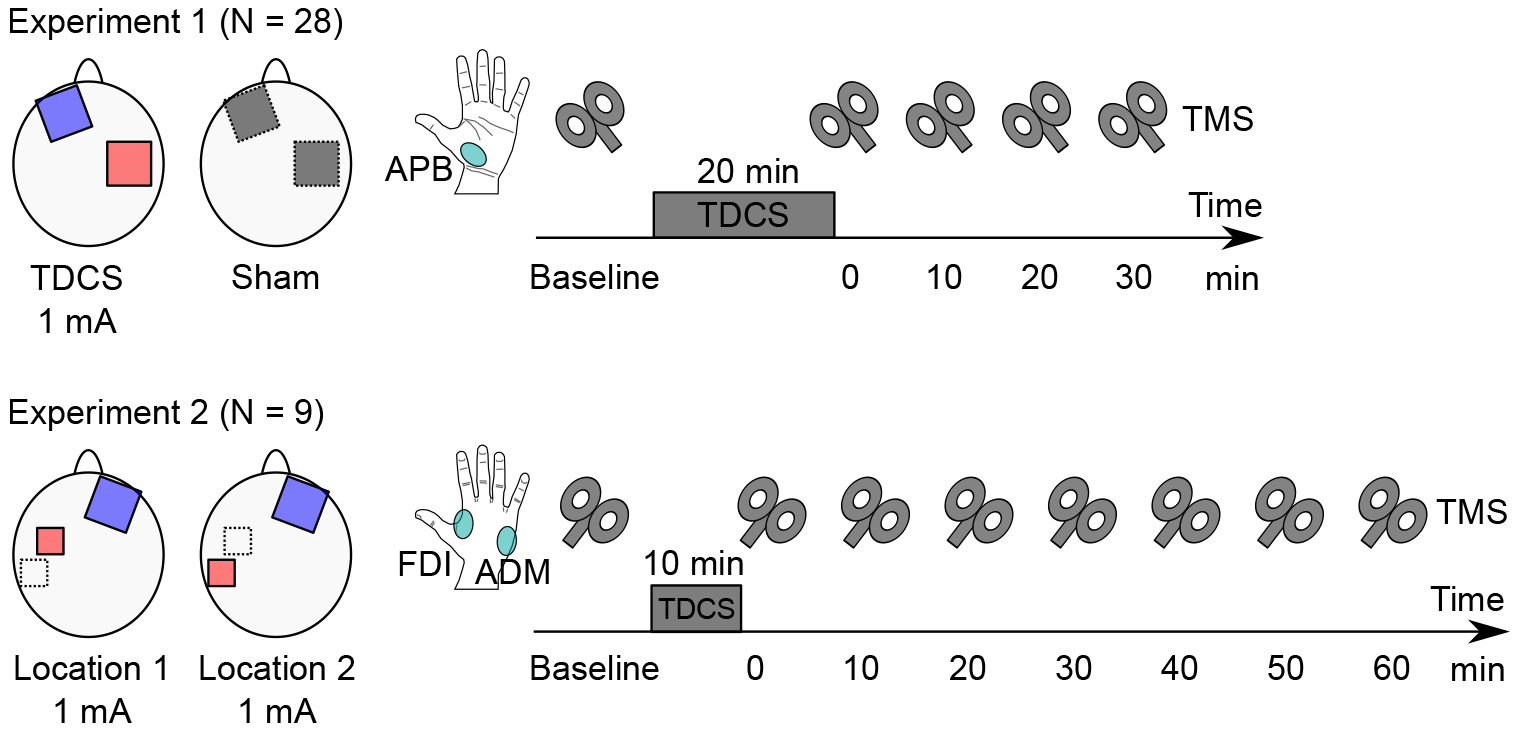
Experimental parameters. In experiment 1, the effect of 20 min anodal TDCS of the right motor cortex was monitored via TMS motor responses in the contralateral APB. In experiment 2, the effect of 10 min anodal TDCS of the left motor cortex was studied in the contralateral FDI and ADM.

#### 2.3.1 Experiment 1

The main experiment employed a double blind, sham-controlled, crossover design to study the effects of anodal TDCS over the right M1 on the MEPs.

The stimulation (anode) electrode (surface area 5×5 cm^2^) was placed over the hand M1 in the right hemisphere. The location of hand M1 (“hand knob” of the precentral gyrus) was identified using an individual T1-weighted MRI and a frameless stereotaxic navigation system (Brainsight 2; Rogue Research, Montreal, Canada). The cathode (surface area 5×5 cm^2^) was placed over the contralateral orbit. Stimulation lasted for 20 min. In the sham condition, current (1 mA) was applied for only the first 15 s. The fade-in/fade-out time was 10 s in each condition.

MEPs of the left abductor pollicis brevis (APB) muscle were recorded before and 0-30 min (with 10 min intervals) after TDCS. During test stimulation, the stimulation intensity with 130% of the RMT was applied 30 times for each time point, and the mean MEP amplitude was calculated.

#### 2.3.2 Experiment 2

An additional experiment was performed to validate the findings of experiment 1 and generalize them to different experimental parameters (opposite hemisphere, shorter duration, and different muscle). Nine subjects participated in this experiment. The effects of anodal TDCS over the left M1 were studied using two different anode locations.

Anode had a surface area of 3×3 cm^2^. Two conditions were studied: The first anode location was above the left hand M1 at the “motor hot spot” where TMS consistently elicited the largest MEPs from the right first dorsal interosseous (FDI) muscle (the navigation system was not available for experiment 2). The second anode location was shifted 2.3±1.6 cm lateral and/or posterior from the hot spot, with the purpose of maximizing the normal component of the EF in hand M1 (Laakso et al., 2016). The choice of the second anode location was guided by EF modelling, and was done individually in each subject. After each condition, a tape measure was used to measure the distances from the centre of the anode to the nasion and both pre-auricular points, and triangulation (Laakso et al., 2018) was used to determine the anode location for subsequent computer modelling. Cathode (surface area 5×5 cm^2^) was placed over the contralateral orbit in both conditions. Stimulation duration was 10 min.

MEPs of the right FDI and abductor digiti minimi (ADM) muscles were simultaneously recorded before and 0-60 min after TDCS. The mean amplitude of sixteen MEP measurements was determined at each time point. The stimulus intensity was set at 130% of the RMT.

### 2.4 Anatomic models and inter-subject registration

T1- and T2-weighted MRI were segmented into distinct tissue compartments. Brain tissues were segmented using the FreeSurfer image analysis software (Dale et al., 1999; Fischl et al., 1999; Fischl and Dale, 2000; Desikan et al., 2006), and remaining tissues were segmented using custom methods implemented in MATLAB (The MathWorks, Inc.). The segmentation process and the tissue conductivities were identical to our previous study (Laakso et al., 2016). The conductivities were (unit: S/m): grey matter 0.2, white matter 0.14, blood 0.7, compact bone 0.008, spongy bone 0.027, CSF 1.8, dura and muscle 0.16, skin and fat 0.08, and eye 1.5.

FreeSurfer with default parameter values was used to generate a mapping from the surface of each individual subject’s brain to that of the standard brain; details of the procedure have been described earlier (Laakso et al., 2016). The standard brain was based on Montreal Neurological Institute (MNI) ICBM 2009a nonlinear asymmetric template (Fonov et al., 2009, 2011).

### 2.5 Electric field modelling

The electrodes were modelled using a two-compartment model consisting of a 1 mm thick rubber pad (0.1 S/m) inserted in a 6 mm thick sponge saturated with physiological saline (1.6 S/m) (Laakso et al., 2016). The electrical sources were a current source (1 mA) and sink (–1 mA) placed inside the rubber pad of the anode and cathode, respectively.

The FEM with cubical 0.5 mm×0.5 mm×0.5 mm first-order elements was used to determine the electric scalar potential *ϕ* from the Laplace-type equation ▽ ·σ▽*ϕ* = 0. The equation was numerically solved using the geometric multigrid method (Laakso and Hirata, 2012) to the relative residual of 10^−6^.

In each subject, the EF was calculated from 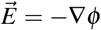 at the depth of 1 mm below the grey matter surface. To separate the EF into three orthogonal components, we calculated the depth vector field 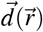 that points from 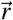 to the nearest point on the inner surface of the skull. In addition, we calculated the outer normal vector 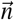 of the grey matter at the hand knob MNI coordinate (40, –20,52) (Yousry et al., 1997; Cárdenas-Morales et al., 2014; Navarro de Lara et al., 2017). 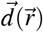 and 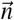 were used to determine the posterior-anterior, medial-lateral, and depth (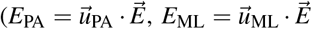, and 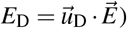) components as follows:

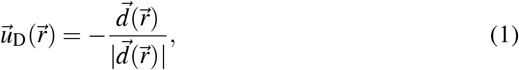

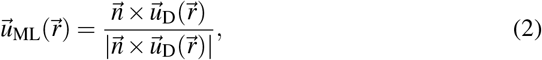

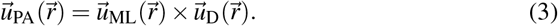

By this definition, *E*_PA_ is perpendicular to the central sulcus at the hand knob, *E*_ML_ is parallel to it, and *E*_D_ points into the brain.

In order to compare the electric fields from different subjects, each component was mapped to the MNI brain (Laakso et al., 2016). The final EFs were represented on a triangular surface mesh of the MNI brain, which consisted of 149319 vertices in the right hemisphere and 148076 vertices in the left hemisphere.

### 2.6 Data analysis

MATLAB was used for all statistical tests. The significance level was P<0.05. All reported P-values are uncorrected for multiple comparisons.

#### 2.6.1 Effects of time and session on MEPs

We first analysed the experimental results without considering the EF, as would conventionally be done in TDCS studies.

Linear mixed effects model was used to study the effects on the MEP amplitude in experiment 1. As fixed effects, we entered the effects of Time (baseline, *t* = 0, 10, 20, and 30 min after stimulation, denoted t0-t30), Session (real TDCS and sham), and their interaction. By-subject intercept and by-subject effect of Session were treated as random effects. P-values of fixed effects were obtained by likelihood ratio tests of the full model versus the model without the effect in question.

For *post hoc* visualization of the effects and their confidence intervals, simpler normalized models were used. In the normalized models, the response was the MEP normalized to the baseline. The effects were otherwise similar to the absolute models, except there was no fixed effect of Session, no random effects, and the intercept was constant 1.

As a measure of overall change in cortical excitability, we calculated the mean MEP amplitude normalized to the baseline over post-stimulation time points (t0-t30)

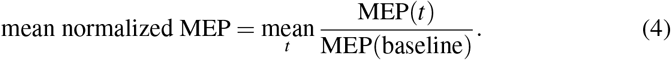

Mean normalized MEPs for the sham and real TDCS were compared using paired t-test, to see whether TDCS had an overall effect at the group level, and the Pearson correlation coefficient, to see whether individuals responded similarly to both sham and real TDCS.

#### 2.6.2 Estimation of important brain regions using PLS regression

We used PLS regression of MATLAB to study whether the measured MEPs in experiment 1 could be explained using the calculated EFs, and, if yes, which brain regions were important for the prediction.

The input data to the PLS regression model were the following. The predictor variables, matrix **X** (28 × 149319), were the EFs (*E*_PA_, *E*_ML_, or *E*_D_) at each vertex on the right hemisphere of the template brain. The dependent variable, vector **Y** (28 × 1), was the mean normalized MEP. The columns of **X** were scaled by dividing them by their sample standard deviations and centred by subtracting their sample mean.

In the initial analysis, the number of PLS components (not to be confused with EF components) was varied from one to three, and the goodness of fit was measured in terms of *R*^2^ (multiple correlation coefficient) and *Q*^2^ (cross-validated *R*^2^). *R*^2^ and *Q*^2^ are the upper and lower bounds, respectively, of how well the model explains the data and predicts new observations (Wold et al., 2001). To calculate *Q*^2^, we used 10-fold cross validation with 1000 Monte-Carlo repetitions. PLS component *i* was defined to be predictively significant if 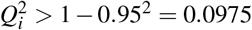 (Abdi, 2010). The analysis was also repeated for the sham MEP data. In this case, the EF should be unrelated to the MEP, and thus, no PLS components should have predictive significance.

After finding the number of significant components, we calculated the variable importance for the projection (VIP) to identify which brain regions were important for predicting MEPs from the EFs. Variables were also selected using competitive adaptive reweighting sampling (CARS) (Li et al., 2009), implemented in the libPLS software library (Li et al., 2018), with 10-fold cross validation and 50 Monte-Carlo sampling runs.

#### 2.6.3 Effect of EF on MEPs

Based on the important variables of PLS regression, we selected a single observation point, 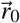, in an anatomically relevant location to better interpret the effect of the EF on the MEP amplitude. This is a valid approach if the effects can be assumed to originate from only one brain region. Unavoidably, the selection was done *post hoc,* and thus, we validated it in another experiment.

At 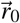, we used a linear mixed effect model to study the potential effect of EF on post-TDCS MEP amplitudes. In addition to the fixed effects of Time, Session and Time×Session, we entered fixed interaction effects of Time×EF and Time×Session×EF for each EF component 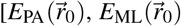 and 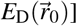 to investigate whether the effect of EF is different between sham and real TDCS. Random effects were by-subject intercept and by-subject effect of Session. Similar to the model without the EF, a normalized model was used for visualization.

Data of experiment 2 were analysed to validate the effects and the choice of 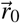 (mirrored to the left hemisphere) found in experiment 1. Linear mixed effects model was used to analyse the fixed effects of Time and EF×Time on the MEP amplitudes. Random effects were by-subject intercept and by-subject effect of Session (anode locations 1 and 2). Note that the model includes both inter- and within-subject effects (two EFs in each subject).

## 3 Results

None of the participants reported side effects.

### 3.1 Overall effect of TDCS on the MEP

The effects were initially analysed without the effect of the EF. The fixed effects of Time (baseline, t0-t30), Session (sham and real) and their interaction on the absolute MEP amplitude were studied using a linear mixed effects model. The model revealed a significant effect of Time (χ^2^ (4) = 17.3, P=0.002). However, the interaction between Time and Session was not significant (χ^2^(4) = 3.049, P=0.5), indicating that both real TDCS and sham changed the MEP amplitudes in a similar manner. Session was not significant (χ^2^(1) = 0.0709, P=0.8), i.e., the baseline MEP amplitudes were not different between real and sham TDCS. Visualization of the normalized MEP (Fig. 2A) showed that the MEP amplitude tended to increase from the baseline for both sham and real TDCS.

To study individual differences in the responses to TDCS, we calculated the mean MEP amplitude normalized to the baseline over all four post-stimulation time points. The group-level, as well as individual data, are presented in Fig. 2B. The mean normalized MEP did not have significantly different group-mean values (paired t-test, *t*(27) = 0.599, P=0.6) nor significant correlation (*R*^2^ = 0.03, P=0.4) between sham and real TDCS.

These results indicated that, while there were no significant differences between sham and real TDCS at the group level, individuals still responded differently to each condition, which is indicated by the lack of significant within-subject correlation between sham and real TDCS. Were these differences due to chance or due to some systematic factor, such as the EF?

**Figure 2:**
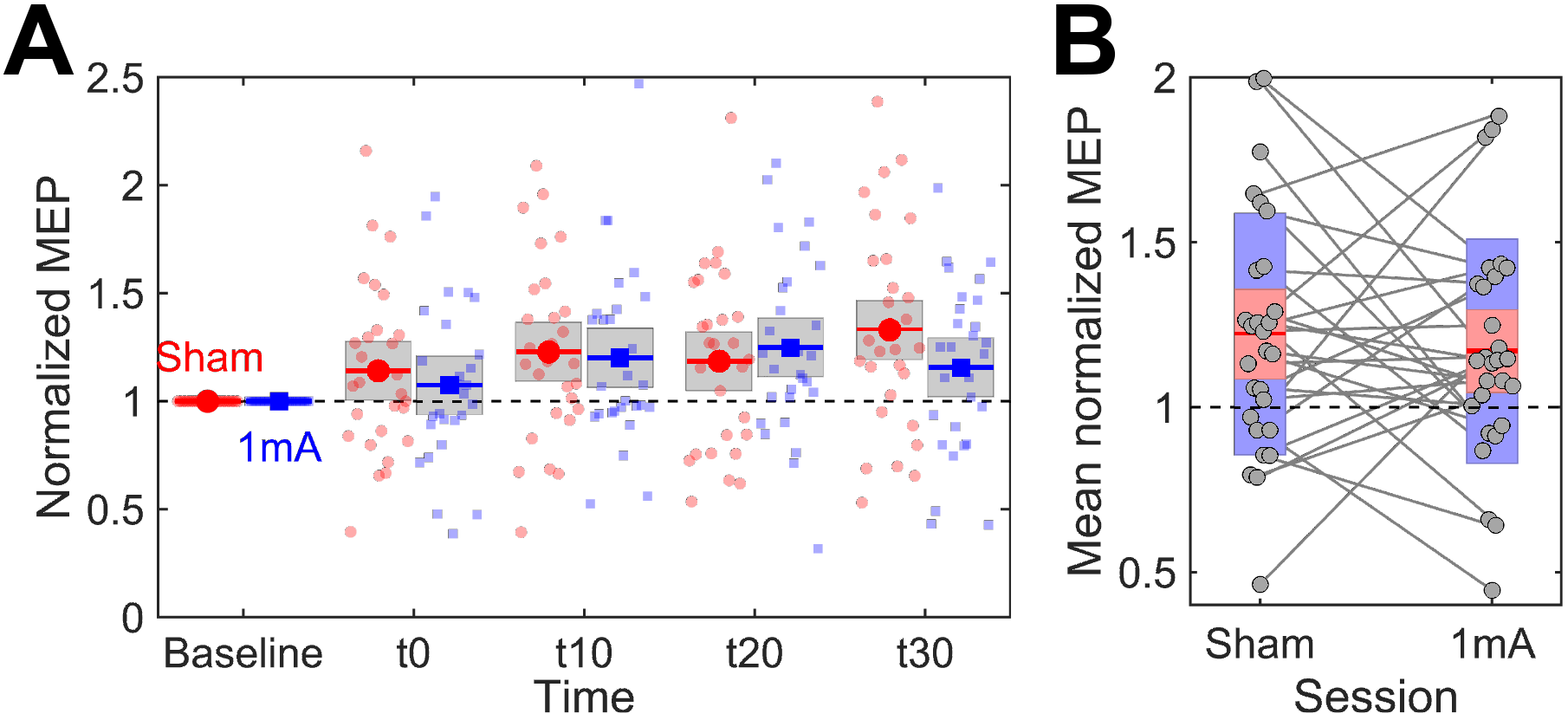
A. Time course of change in the normalized MEP amplitude (N=28). Markers are predicted values and bars are 95% confidence intervals from the linear model. Small markers are individual observations. B. Grand mean normalized MEP over post-stimulation time points. Circles represent the data for individual subjects. Mean value is indicated by the horizontal line, and coloured bars represent the standard deviation (light blue) and 95% confidence interval (light red).

### 3.2 Calculated EFs and PLS regression

The EFs were calculated individually in 28 subjects participating in experiment 1 and separated into PA, ML, and depth (*E*_PA_, *E*_ML_, and *E*_D_) components. Each component was then registered to the standard brain for further analysis. Figure 3 shows the average EF components in the right hemisphere. The EF in hand M1 is primarily in the depth direction. The *E*_PA_ component is smaller than the *E*_D_ component in hand M1, and its maximum is found in the frontal areas. The *E*_ML_ component is small in hand M1.

**Figure 3:**
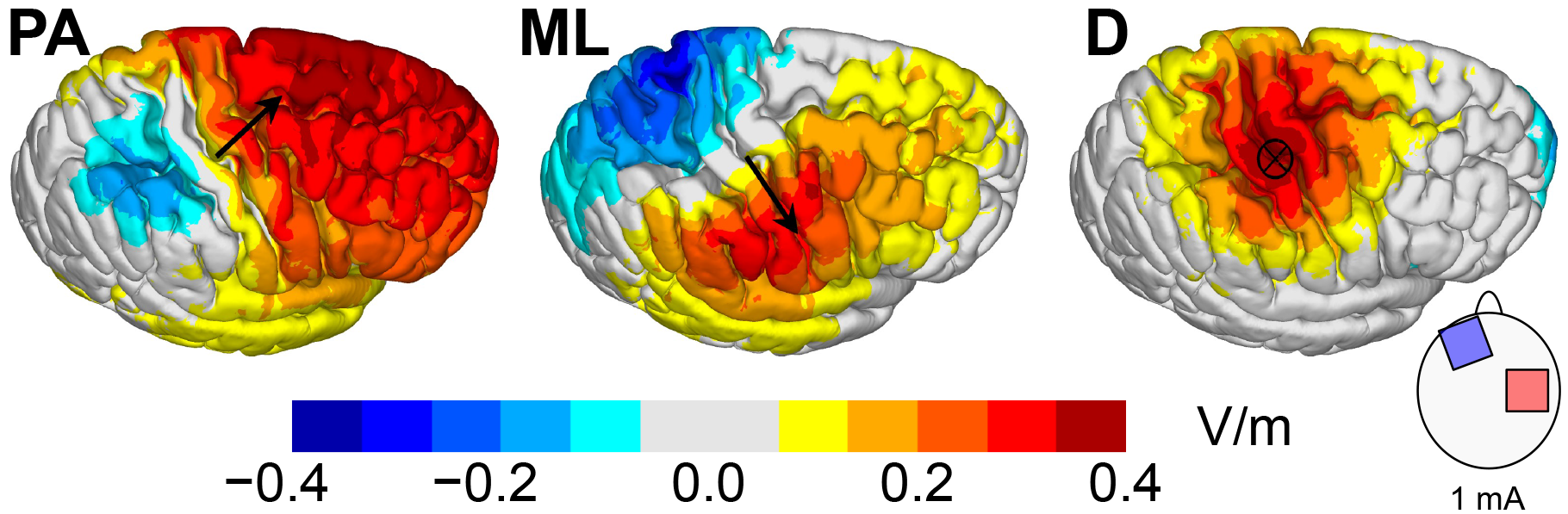
Group-average EF (N=28) separated into three orthogonal components: posterior-anterior (PA), medial-lateral (ML) and depth (D). The electric fields were first divided into components in each subject, after which the components were registered to a common template and finally averaged. Arrows visualize the component directions in the hand motor area.

The initial PLS regression analysis with *E*_D_ as the predictor gave one predictively significant PLS component (*R*^2^ = 0.77, *Q*^2^ = 0.18, Table 1). Using other EF components (*E*_PA_ or *E*_ML_) as predictors or sham MEPs as dependent variables did not result in any predictively significant PLS components (Table 1). The model with one PLS component and *E*_D_ as the predictor was used for the subsequent analysis. PLS score plots indicated no violations of homogeneity or curvature of the data. Normal probability plots were used to verify the normality of residuals, and no clear outliers were detected in residual plots.

**Table 1:** Explained 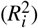 and predicted 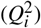 variance of the first three PLS components (*i*). The predictor variables are the EF components and the dependent variables are the mean normalized MEP of either real TDCS or sham stimulation.

| Predictor | $R_1^2$ | $R_2^2$ | $R_3^2$ | $Q_1^2$ | $Q_2^2$ | $Q_3^2$ |
| --- | --- | --- | --- | --- | --- | --- |
| *Real TDCS* |  |  |  |  |  |  |
| $E_{PA}$ | 0.71 | 0.24 | 0.04 | -0.07 | -2.49 | -18.38 |
| $E_{ML}$ | 0.89 | 0.08 | 0.03 | -0.55 | -10.00 | -35.66 |
| $E_D$ | 0.77 | 0.22 | 0.01 | 0.18* | -2.62 | -126.18 |
| *Sham* |  |  |  |  |  |  |
| $E_{PA}$ | 0.75 | 0.20 | 0.03 | -0.63 | -4.60 | -32.47 |
| $E_{ML}$ | 0.56 | 0.40 | 0.02 | -0.54 | -2.12 | -33.64 |
| $E_D$ | 0.87 | 0.12 | 0.02 | -0.21 | -7.77 | -63.31 |

\* $Q^2 > 0.0975$

Next, we investigated which brain regions were important for predicting the MEP from *E*_D_. Figure 4 shows the results obtained using the VIP and CARS methods. The regions with high VIP in Fig. 4A and the clusters in Fig. 4B are candidates for the site of action where the EF has an effect on the MEP. However, the number of candidate sites is relatively large, which is due to the fact the PLS regression used no information about the functional relevance of each brain region or the EF strength (EFs were normalized before they were input into the model).

**Figure 4:**
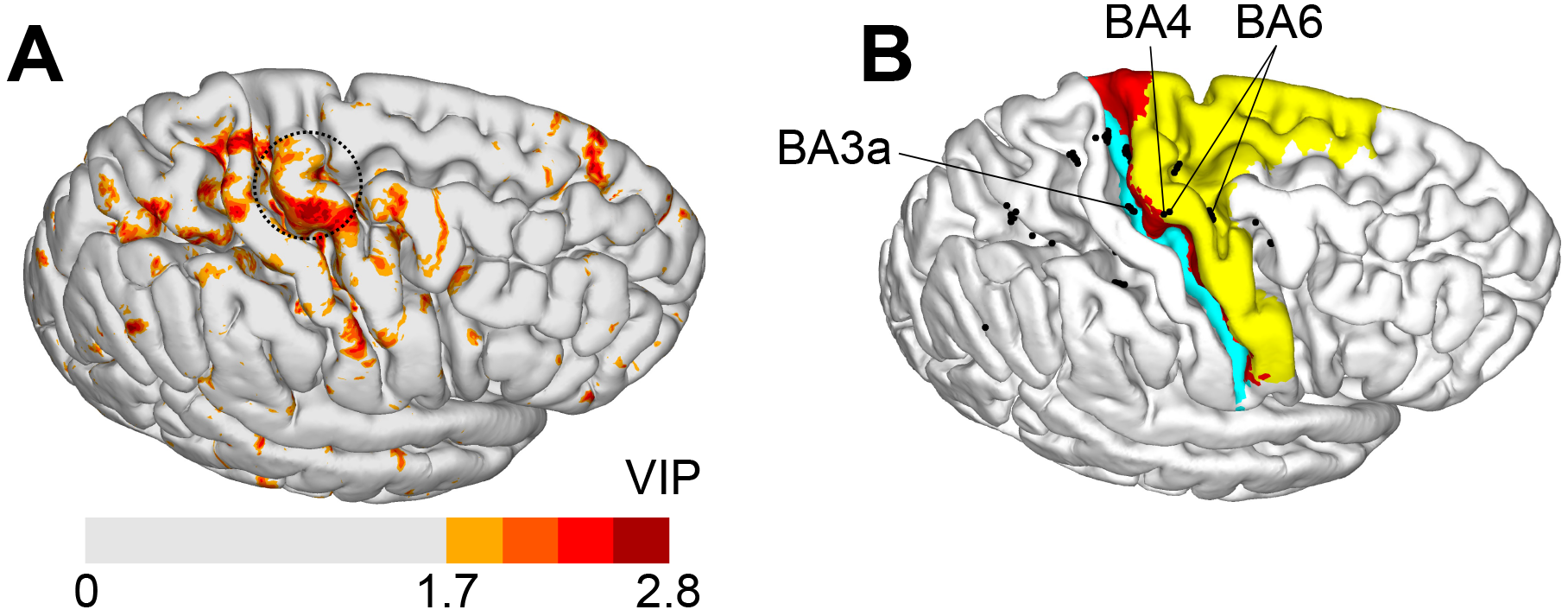
Important brain regions estimated using PLS regression analysis. A. VIP, indicating the relative importance of brain regions for predicting post-TDCS MEP from the EF. Coloured regions have VIP higher than the 90th percentile. Circle indicates the inverted omega of the hand knob. B. Important variables selected by CARS, indicated by black dots. Colours are Brodmann areas 3a, 4, and 6, which have been estimated using the probabilistic cytoarchitectonic map of FreeSurfer. Potentially relevant variables are labelled by their estimated Brodmann areas.

To narrow down the number of candidate sites, we used the data on EF strength (Fig. 3), anatomical location of the hand area (Fig. 4A), and probabilistic cytoarchi-tectonic atlas (Fig. 4B). This reduced the likely sites of action to the anterior subarea of BA4 or associated areas (BA3a or BA6) near the hand knob (labelled in Fig. 4B). For the subsequent analysis, we selected an observation point at the border of BA4a and BA6, labelled “BA4” in Fig. 4B. In MNI coordinates, the observation point was 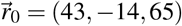.

The choice of 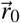 from among several candidates is motivated by the fact that 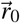 is close to the TMS hotspot of the ABP muscle projected to the cortex, (–41±4,–16±4.60±4), measured in the left hemisphere (Diekhoff et al., 2011). The observation point is also close to the projected TMS hotspots of the FDI muscle, (–43±7,–10±8.61±6) (Volz et al., 2015) and (–37±3,–19±3,66±2) (Bungert et al., 2016). Next, we investigated how the EFs at 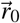 affect the MEPs and their time course. Finally, we performed an additional experiment to validate the choice of 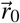.

### 3.3 Effect of EF in right M1

The EFs calculated at 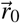 were input into a linear mixed effects model as fixed effects. Based on the results of the PLS regression, we considered only the *E*_D_ component. Adding the *E*_PA_ and *E*_ML_ components did not improve the linear mixed effects model significantly (likelihood ratio test of full model versus model without the terms including *E*_PA_ and *E*_ML_ components, χ^2^(16) = 14.961, P=0.52). The summary statistics of 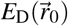 were mean±SD: 0.38±0.11 V/m and range: 0.20-0.63 V/m.

Likelihood ratio test showed that the three-way interaction effect 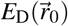×Time× Session was significant (χ^2^(4) = 15.618, P=0.0036), indicating that the EF had a differential effect depending on the time point and whether sham or real TDCS was used. 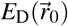×Time interaction was not significant (χ^2^(4) = 3.4296, P=0.49), i.e., the EF did not have significant time-dependent effects that were common to both sham and real TDCS. Similarly to the model without the EF (Sec. 3.1), the effect of Time was significant (χ^2^(4) = 18.704, P=0.00089). Session as well as Session×Time were not significant (P>0.5).

Normalized MEPs were used to visualize the identified effects (Fig. 5). In the case of real TDCS, but not sham, the normalized MEPs were modulated by the individual EF, which explains the effect of 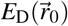×Time×Session. In subjects with less positive 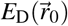, real TDCS tended to increase the MEP compared to the baseline, whereas in subjects with a more positive 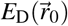, the MEP slightly decreased or did not change compared to the baseline. The effect persisted over all time points. For sham stimulation, 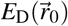 did not affect the MEP at any time point.

**Figure 5:**
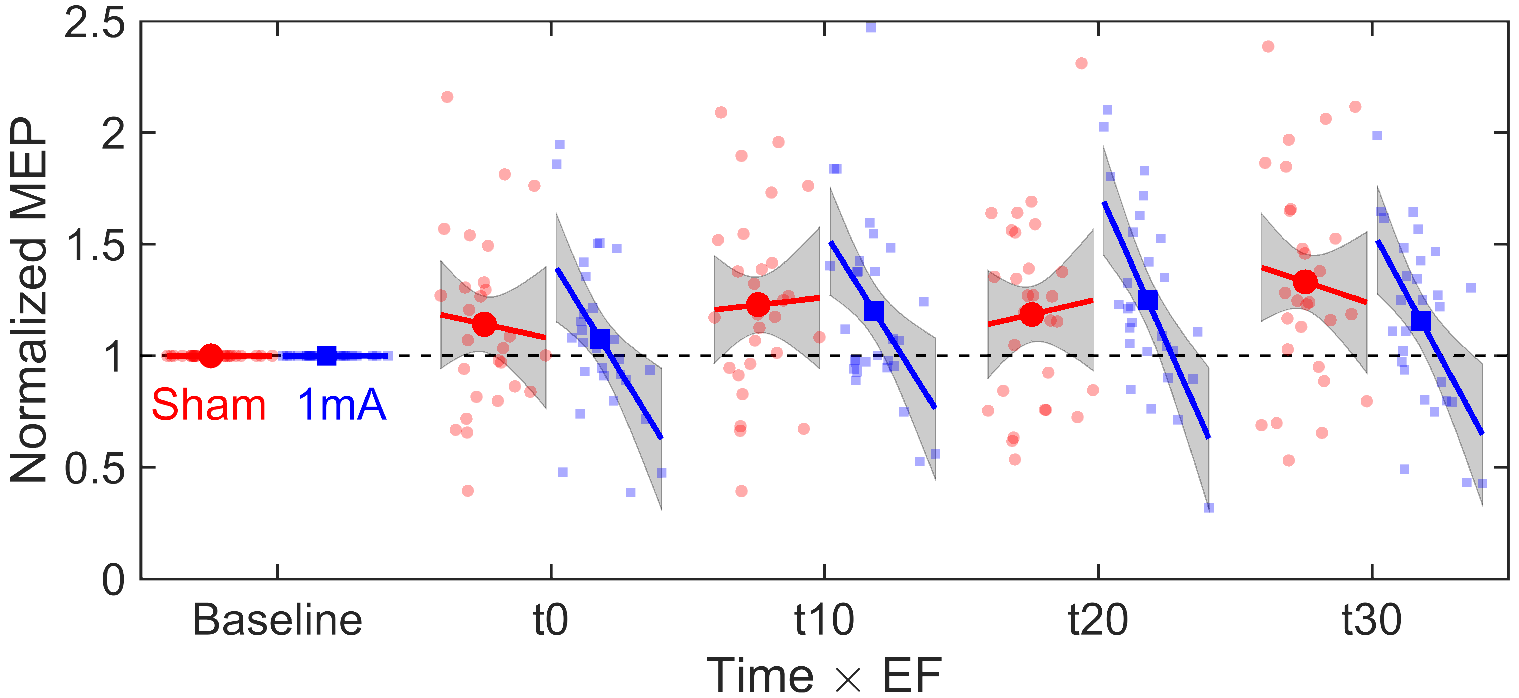
Effect of the EF depth component and time on normalized MEP for 20 min 1 mA anodal TDCS of the right M1 (N=28). MEPs were measured in the left APB muscle. Markers show the mean values. Lines and shaded areas are the regression lines and the 95% confidence intervals from the linear model (range: 0.20-0.63 V/m).

Because the effect of 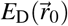 on the normalized MEP did not differ significantly between post-stimulation time points (Fig. 5), we used simple linear regression to characterize the effect of 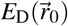 on the mean normalized MEP. The fitted linear model (*R*^2^ = 0.42, P=0.00019) was

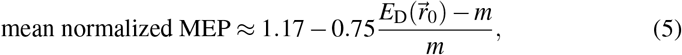

where *m* = 0.38 V/m is the sample mean of 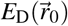. The 95% confidence intervals of the intercept and slope are [1.07,1.27] and [−1.11,−0.40], respectively.

### 3.4 Effect of EF in left M1

An additional experiment with nine subjects was performed to validate the choice of 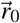 and study whether the findings of experiment 1 are valid for the opposite hemisphere and different stimulation duration.

Figure 6 illustrates the EFs of both electrode configurations used in experiment 2. On average, anode location 2 produced larger *E*_PA_ and smaller *E*_D_ in hand M1 than anode location 1. The EFs are slightly more localized than those in experiment 1 (Fig. 3) owing to the smaller anode surface area.

The EFs were determined at 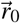 (with mirrored *x* coordinate), and linear mixed effects model was used to analyse the fixed effects of Time and 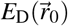×Time on the MEPs recorded in the FDI and ADM muscles. Again, we report the results only for the model with the *E*_D_ component; adding the *E*_PA_ and *E*_ML_ components did not improve the model significantly (FDI muscle: χ^2^(14) = 5.7521, P=0.97; ADM muscle: χ^2^(14) = 20.938, P=0.10). The summary statistics of 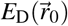 were mean±SD: 0.44±0.17 V/m and range: 0.09-0.76 V/m.

**Figure 6:**
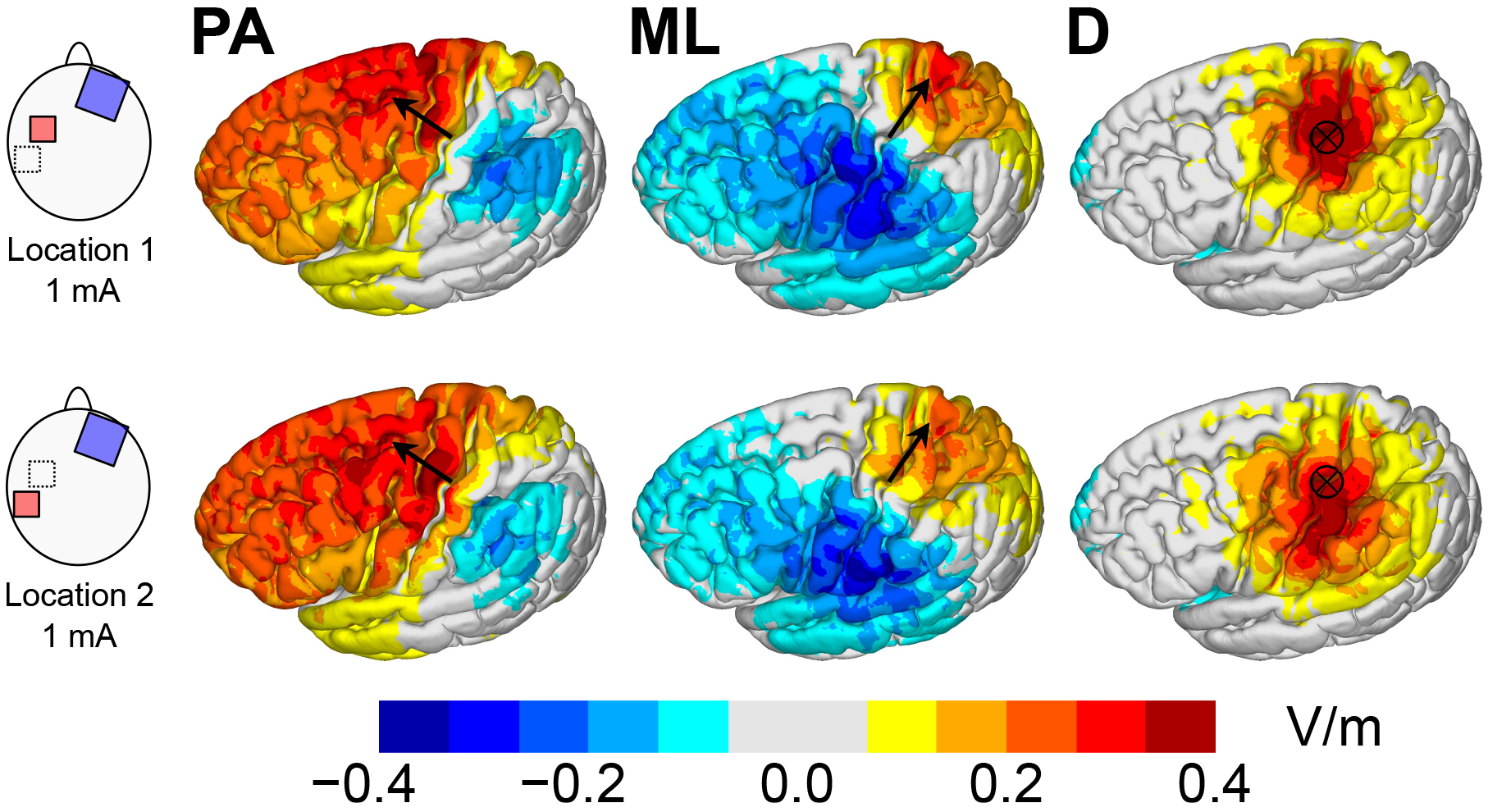
Group-average EFs (N=9) for two electrode configurations of experiment 2.

For the FDI muscle, likelihood ratio tests showed that the interaction term 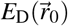×Time was significant (χ^2^(7) = 24.447, P=0.00095), indicating that the change in the MEP with time depended on the individual EF. Time was not significant (χ^2^(7) = 4.1397, P=0.76), showing that the average MEP at the group level did not change significantly from the baseline. For the ADM muscle, neither 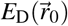×Time nor Time were significant (P>0.18).

Normalized MEPs were used to visualize the results for the FDI muscle. Figure 7 shows that more positive 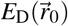 increased the MEP, whereas less positive 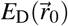 decreased the MEP compared to the baseline. Interestingly, the direction of the effect was opposite compared to experiment 1. In addition, the effect persisted at least 60 min after stimulation, which was the last time point measured.

Figure 7 also illustrates the consistency of the EF effect within individual subjects. Immediately after stimulation (t0), eight out of nine subjects showed an increased MEP in the FDI muscle for an increased EF (P=0.020, from binomial distribution), consistently with the group-level effect of EF. The within-subject effect was not significant for later time points (P>0.24). We note that, because we mainly considered the normal component (approximately *E*_PA_) when selecting the second anode location, some subjects had very similar 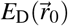 in both conditions, and one subject had a higher 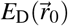 for anode location 2. Interestingly, the three subjects with the largest absolute change in 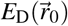 between the anode locations had a consistently positive EF-MEP relationship at all but one time point (one subject at t40).

Simple linear regression gave the following formula for estimating the mean normalized MEP (FDI muscle, average over time points t0-t30 for consistency with experiment 1) from *E*_D_ (*R*^2^ = 0.48, P=0.0015):

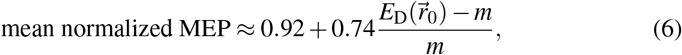

where *m* = 0.38 V/m (chosen to be the same as in (5)). The 95% confidence intervals of the intercept and slope are [0.74,1.11] and [0.33,1.15], respectively. Compared to experiment 1, the effect of EF is of similar size but in the opposite direction, and the intercept is smaller.

**Figure 7:**
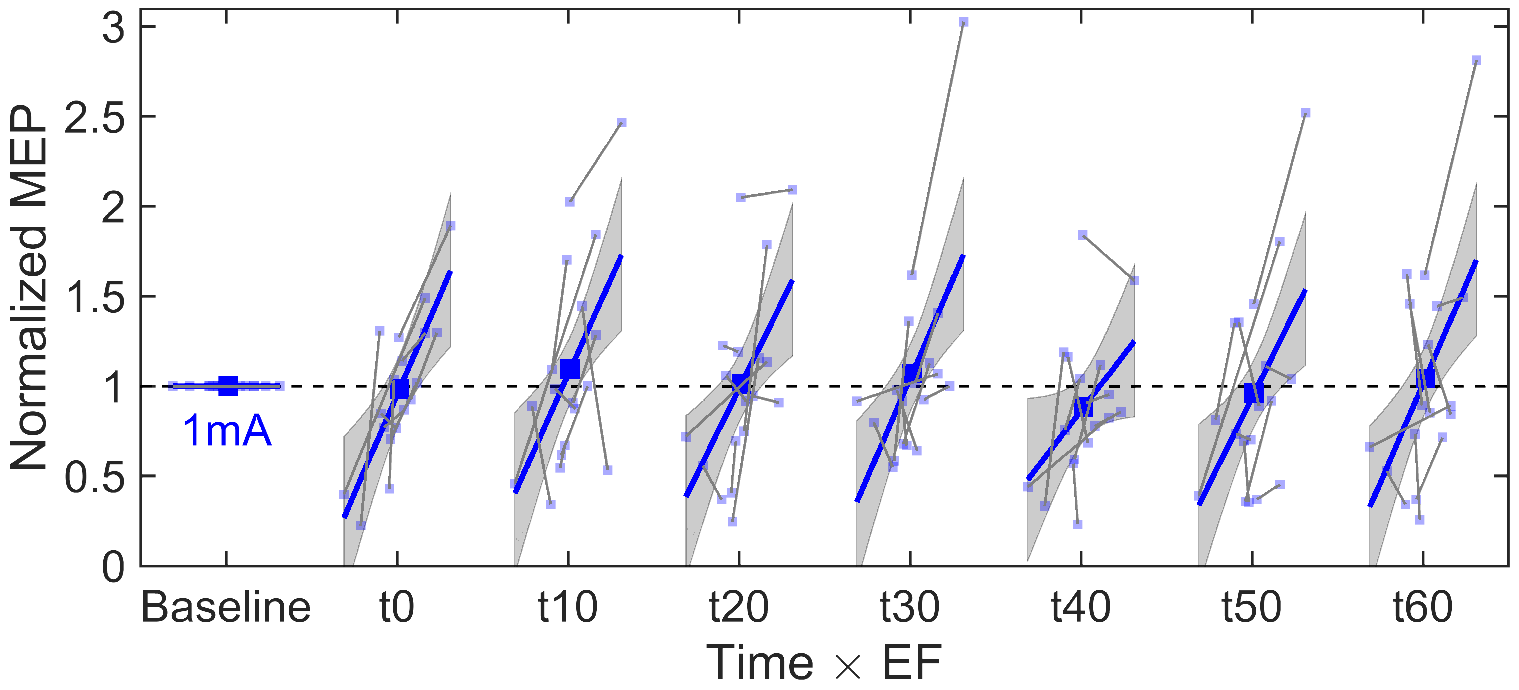
Effect of the EF depth component and time on normalized MEP for 10 min 1 mA anodal TDCS of the left M1 (N=9). MEPs were measured in the right FDI muscle. Each subject was stimulated at two different electrode locations. Large markers show the mean values and lines and shaded areas are the regression lines and the 95% confidence intervals from the linear model (range: 0.09-0.76 V/m). Small markers and grey line segments show the observations from individual subjects.

## 4 Discussion

The main finding of this study is that subjects responded differently to TDCS depending on the individually modelled EFs. The findings are important for the inter-individual variability and dosing of TDCS.

### 4.1 Inter-individual variability

Each individual’s response to TDCS depended on the individual EF. The effect of EF was strong and persistent, explaining approximately 40-50% of inter-subject variance in mean normalized MEPs, and lasting at least 60 min post-stimulation. This suggests that inter-individual differences in the EF dose are essential for understanding the effects of TDCS.

Existing TDCS protocols that have been used since the early 2000s typically apply the same input current to all subjects (Nitsche et al., 2008; Horvath et al., 2015; Woods et al., 2016). Based on our findings, this approach is problematic: Subjects with the lowest and highest EF strengths may respond oppositely to the same input current, which could result in weak or not significant findings at the group level. Indeed, in both of our experiments, which employed fairly typical TDCS parameters, no significant group-level differences compared to sham or baseline would have been found without considering the EF. For comparison, several previous studies have also reported small group-level responses but high inter-individual variability (Wiethoff et al., 2014; López-Alonso et al., 2014; Chew et al., 2015).

In previous studies, variability of individuals’ responses to TDCS have been characterized using clustering analysis (Wiethoff et al., 2014; Chew et al., 2015; Strube et al., 2016; Ammann et al., 2017). A simple, but not always appropriate (Ammann et al., 2017), way to study the response profile is to group the subjects by whether the mean normalized MEP increased or decreased compared to the baseline. Using this approach, Wiethoff et al. (2014) found that 74% subjects had “facilitatory” response for anodal TDCS (left M1, 2 mA, 10 min, N=53). In the study of Strube et al. (2016), the percentage of “facilitatory” responders was 61% (leftM1,1 mA, 13 min, N=59). Using the same criteria and ignoring sham, our results would have shown that 75% had “facilitatory” and 25% “inhibitory” response (see Fig. 2B). Due to the relationship between the EF and MEP, the mean 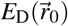 of the groups would have been significantly different, 0.35 ± 0.09 and 0.48 ± 0.11 V/m, respectively (t-test, *t*(26) = 3.2064, P=0.0035). This illustrates that groups of responders and non-responders found in previous works may have been due to differences in the EF.

If the EFs are indeed related to TDCS after-effects, the EF dose should be considered in TDCS protocols. Our initial results in experiment 2 suggest that altering the EF can change the MEP in the expected way, at least immediately after stimulation and for large changes in EFs, suggesting that EF models could be used to individually control TDCS, and hence, reduce variability. However, this needs to be confirmed in additional studies.

### 4.2 Non-linear effects

The effect of EF depended on the stimulation parameters, indicating possible nonlinearity. In our first experiment with 20 min 1 mA anodal TDCS of the right M1 and MEP recordings in the contralateral APB muscle, a more positive *E*_D_ decreased the MEP. The effect was equally strong but opposite in the second experiment, where we used 10 min 1 mA anodal TDCS of the left M1 and measured the MEP in the contralateral FDI muscle (the effect was not significant in the ADM muscle). This inconsistency may be due to differences in the stimulation parameters, particularly the stimulation duration.

Previous studies have shown that the stimulation duration has a strong effect on the responses. Classically, longer durations are thought to produce stronger effects (Nitsche et al., 2008), as reported by Nitsche and Paulus (2000), who applied 1-5 min of anodal 1 mA current (N=12). However, for longer stimuli, the effects are nonlinear. Monte-Silva et al. (2013) found that 13 min of 1 mA anodal TDCS of left M1 (N=15) enhanced motor cortical excitability, measured by MEP amplitudes in the contralateral ADM muscle. Importantly, doubling the duration to 26 min reversed the effect, reducing the excitability (Monte-Silva et al., 2013). Diminished excitability has also been found using nearly identical parameters with our study (right M1, 1 mA, 20 min, FDI muscle, N=11) (Simis et al., 2013), whereas anodal stimulation for 9–13 min typically enhances excitability (Nitsche et al., 2008). We hypothesize that the opposite effect of EF for 10 min and 20 min stimulation found in the present study and the previous group-level findings (Monte-Silva et al., 2013; Simis et al., 2013) are in fact different manifestations of the same non-linear phenomena.

In addition to duration, the effects of TDCS are known to be non-linear dependent on the current intensity (Batsikadze et al., 2013; Kidgell et al., 2013; Jamil et al., 2017). The non-linear group-level effects of current intensity may be related to our finding of opposite effects in low-EF and high-EF subjects. Therefore, understanding the nonlinearity with both duration and magnitude is essential if EF models are used to control TDCS.

### 4.3 Which sites are affected by EF?

For the APB and FDI muscles, we found that the calculated EFs in the anterior bank of the central sulcus in the lateral part of the hand knob [MNI coordinate (±43,–14,65)] were significantly related to the measured MEPs. Cytoarchitectorally, the site is at the border between the primary motor cortex (BA4a) and premotor areas (BA6). The effects failed to reach significance for the ADM muscle, which indicates that the EF at a different, perhaps more medial, site should have been considered for modelling the MEPs of the ADM muscle.

Based on our findings, an attractive hypothesis is that the after-effects of TDCS are mediated by the local EF in hand M1. This is plausible also because long-lasting effects of weak EFs in M1 have been reported *in vitro* (Bikson et al., 2004; Fritsch et al., 2010). Because the site is located very close to the TMS hotspot of the muscle in question (Diekhoff et al., 2011; Volz et al., 2015; Bungert et al., 2016; Laakso et al., 2018), the EF might change the excitability of the same site that is activated by TMS. Previously, Fischer et al. (2017) argued that the local effect of EF on M1 was unlikely, and the effects more likely originated from regions outside M1. Their arguments based on the finding that a multifocal TDCS montage that produced weaker EF in left M1 resulted in a greater increase in excitability than conventional TDCS (2 mA, 10 min, FDI muscle, N=15) (Fischer et al., 2017). Our findings show that we cannot assume a positive EF strength-response relationship in M1, and thus, the findings of Fischer et al. (2017) may have actually been due to a local effect of EF in M1.

We also found that the direction of the EF in hand M1 is important for explaining the changes in the MEPs. After separating the EF into three orthogonal components, we found that only the *E*_D_ component had a significant effect on the MEPs. This was the case also in experiment 2, where we aimed to manipulate the *E*_PA_ component by shifting the anode location. At the anterior bank of central sulcus, the positive direction of *E*_D_ is “into” the brain. Previously, Rawji et al. (2018) studied the excitability changes using two electrode montages that produced EFs approximately in the PA and ML directions (left M1, 1 mA, 10 min, FDI muscle, N=15); the depth component was not reported. They found that the EF in the PA direction modulated the MEPs, decreasing the excitability, while the EF in the ML direction did not (Rawji et al., 2018). The results of experiment 2 are in line with these findings, except that the effective component was *E*_D_, not *E*_PA_. Namely, moving the anode posteriorly, which on average produced smaller *E*_D_ and at the same time larger *E*_PA_, decreased the excitability.

### 4.4 Limitations

Unexpectedly, sham stimulation significantly increased the group-average MEP in experiment 1, but the increase was unrelated to the EF. We are unsure of the reasons, as other recent sham-controlled studies have not shown any significant effects of sham (Jamil et al., 2017; Ammann et al., 2017; Rawji et al., 2018). The effect of sham persisted at least 30 min after stimulation, which highlights the importance of sham control in TDCS studies.

A significant EF-MEP relationship does not imply a causal effect. The true physical agent of TDCS may as well be another electrical or electrochemical quantity that is proportional to the EF. For instance, the physical agent could be the total movement of charge through unit area, which depends monotonically on both the EF and stimulation duration and is particularly interesting because it could be used to combine the effects of both duration and intensity. We also note that the modelled EFs are approximative due to uncertainty in the biophysical properties of tissue *in vivo* (Opitz et al., 2017).

Our results showed that the EF-MEP relationship is significant in at least one site in M1. However, this does not necessarily mean that it is the true site of action. As the EFs in the cortex are spatially diffuse and highly collinear, any closely adjacent site or even some distant sites would produce a very similar relationship between the EF and MEP. The PLS regression analysis revealed many other regions that might potentionally be important for TDCS after-effects, which, however, seems less likely than the local effect of EF in M1. Finally and obviously, our approach fails to find any excitability changes that are invisible in the MEP recordings.

Despite non-linearity discussed earlier, the EF-MEP relationship was locally linear in the studied conditions, and thus, the effects could be studied using linear models. Because the linear approximation does not hold globally, our findings should not be extrapolated to other experimental conditions.

### 4.5 Conclusion

The after-effects of motor cortical TDCS are modulated by the EF or a related electrical quantity. Considerable part of variability commonly observed in TDCS studies may be due to inter-individual differences in EFs. The EF probably acts locally in M1, and the effect depends on the field direction. Individual EF models could be used to reduce inter-subject variability and control the effects. However, this requires understanding of the non-linear effects of EF.

## Acknowledgements

S.T. was supported by KAKENHI (No. 17H00869 and 16H03201).

